# Autophagy-Mediated Antitumor Effects of BCG and Imiquimod in Oral Squamous Cell Carcinoma

**DOI:** 10.64898/2026.07.30.741756

**Authors:** Nancy A. Abdelmoneim, Magi N. Moussa, Alaa A. Elmorsy, Manal I. Elnouaem, Omneya R. Ramadan, Hend M. Abdelhamid, Radwa A. Mehanna, Ashraf K. Awaad, Enas M. Omar, Marwa M. Afifi

## Abstract

Autophagy and Toll-like receptor (TLR) signaling are both implicated in cancer progression, but whether they act as tumor suppressors or promoters remains unclear. To explore this relationship, we investigated the role of autophagy downstream of therapeutic TLR activation in oral squamous cell carcinoma (OSCC). Using the FDA-approved TLR agonists Bacillus Calmette-Guérin (TLR2/4) and imiquimod (TLR7), we assessed their effects in vitro on SCC-4 cells and in vivo in a chemically induced OSCC hamster model. Both agents, alone or in combination with each other or with Monophosphoryl Lipid-A induced robust autophagy, as measured by LC3B staining in vitro and flow cytometry in vivo. Autophagy induction correlated with reduced tumor volume and prolonged survival, with outcomes comparable to cisplatin, the current standard chemotherapeutic, but with less treatment-associated morbidity and mortality. Autophagy was also associated with cisplatin antitumor effects. Notably, imiquimod produced the most pronounced and sustained autophagic and antitumor effects. To our knowledge, this is the first study to directly link the therapeutic efficacy of TLR agonists in OSCC to autophagy modulation, providing both mechanistic and translational insights into their potential as immunotherapeutic agents.

## Introduction

Oral squamous cell carcinoma (OSCC) accounts for over 90% of lip and oral cavity cancers and remains a major public health burden, with an estimated 5-year survival rate below 50% despite advances in surgery, radiotherapy, and chemotherapy [1–3]. In Egypt, the mortality-to-incidence ratio approaches 63%, reflecting limited therapeutic success [1]. These poor outcomes underscore the need for novel therapeutic strategies that not only target tumor cells directly but also engage the host immune system to achieve durable control. Immunotherapy has transformed the management of several cancers, yet its clinical impact on OSCC remains modest, in part due to tumor immune evasion and the lack of optimized immunomodulatory agents tailored for this disease [4,5].

Toll-like receptors (TLRs) are pattern-recognition receptors (PRRs) expressed on immune and non-immune cells, including cancer cells, and play a central role in bridging innate and adaptive immunity. TLR agonists such as Bacillus Calmette-Guérin (BCG; TLR2/4), Imiquimod (TLR7), and Monophosphoryl lipid A (MPLA; TLR4) are FDA-approved for cancer therapy, either as stand-alone agents or as vaccine adjuvants [6–8]. Upon activation, TLRs initiate inflammatory cascades that promote dendritic cell maturation, cytokine secretion, and tumor-specific immune responses.[6] However, TLR signaling in cancer is context-dependent: while it can stimulate anti-tumor immunity, persistent or dysregulated activation may instead promote immune evasion, angiogenesis, and metastasis [9–11]. Identifying strategies that bias TLR responses toward tumor suppression is therefore critical for effective therapeutic application.

Autophagy, a lysosome-dependent degradation pathway, is one downstream effector of TLR signaling [12–14]. Under basal conditions, autophagy maintains cellular homeostasis by recycling cytoplasmic components; it is upregulated in response to stress [15–17].

While initially cytoprotective, sustained autophagy can culminate in caspase-independent programmed cell death, termed autophagy-dependent cell death (ADCD), characterized by extensive autophagosome formation and cytoplasmic vacuolization without chromatin condensation [18,19]. Like TLR signaling, autophagy can be tumor-suppressive or tumor-promoting depending on the context [20], suggesting that deliberate modulation of autophagy could influence the therapeutic outcome of TLR-based interventions.

Although BCG and imiquimod have been evaluated for their immunomodulatory effects in OSCC, and other head and neck cancers, their ability to induce autophagy, and autophagy-mediated cell death in this malignancy has not been systematically investigated. Prior reports have hinted at potential anti-tumor benefits, but these were limited to small studies or historical trial with inconsistent outcomes [21,22,31,23–30]. A recent pilot clinical trial explored imiquimod as a neoadjuvant to surgery in OSCC, emphasizing its safety and promise [32]. Autophagy is a possible mediator of imiquimod action [33], however, mechanistic insights into its role in OSCC are lacking. Furthermore, the comparative effects of multiple clinically relevant TLR agonists, including MPLA, on both tumor cell-intrinsic and immune-mediated pathways in OSCC remain unknown. In this study, we directly compare BCG, imiquimod, and MPLA for their capacity to activate distinct TLR pathways, alter receptor localization, and trigger autophagy in OSCC models. By integrating *in vitro* mechanistic assays with *in vivo* tumor models, we identify conditions under which TLR activation can be steered toward a cytotoxic autophagic outcome, thereby uncovering a previously unexplored therapeutic avenue for TLR-based immunotherapy in OSCC.

## Materials and methods

### Cell Line

The *In-vitro* part of the study was conducted at the Center of Excellence for Research in Regenerative Medicine and its Applications (CERRMA), Faculty of Medicine, Alexandria University. We used American Type Culture Collection (ATCC) authenticated human oral squamous cell carcinoma cancer cell line SCC-4 (ATCC^®^ CRL1624™).

### Cytotoxicity Assay for TLRs 2,4, and 7 Agonists

SCC-4 cells were seeded in 96 well plates at the density of (5 x 10^3^ cells/100 µl) in each well. Plates were incubated in 5% CO2, 37 °C for 24 hours. The cells were then incubated with eleven concentrations of BCG (Immune BCG-T^®^, VACSERA, Egypt) (0.5, 1, 1.5, 2, 2.5, 3, 3.5, 4, 4.5, 5 and 5.5 mg/ml), twelve concentrations of Imiquimod (Cat. No. 99010-64-7, Reference Standard 1338313 USP) (5, 20, 30, 40, 50, 60, 70, 80, 90, 100, 110 and 120 µg/ml), and five concentrations of MPLA (Cat. No. ALX-581-205-C100, ENZO life sciences) (1,2,3,4, and 5 µg/ml) for 48 hrs. Details of drug preparation are found in supplementary material. Negative control wells received no treatment. Cell counting kit- 8 (CCK-8) reagent (Dojindo molecular technologies inc, CA, USA, Cat. No. CK04-11) was applied as 10% of the total volume of media to all wells and incubated for 4 hrs. in 5% CO2, at 37 °C. Then, the color absorbance was read at 450 nm using an automated microplate reader (BioTek® Instruments, VT, USA); All values were corrected with the reference wavelength at 630 nm and normalized against the mean value of the negative control wells. The half maximal inhibitory concentration (IC50) and IC80 for BCG were interpolated using a fit spline curve. For imiquimod, none of the tested doses reached IC50 at 48 hours; therefore, IC80 values were interpolated, and IC50 was subsequently estimated using non-linear regression analysis (GraphPad Prism version 9, GraphPad Software Inc., San Diego, CA).

### Verification of TLR 4 and 7 Expression Upon the Application of BCG and Imiquimod by Confocal Laser Scanning Microscopy

Six-well plates were prepared with a seeding density of 80,000 cells/well. A cover slip was put in each well. When cell reached confluence, two doses of each drug were selected and applied, 0.5 and 5 mg/ml for BCG, 5 and 120 µg/ml for Imiquimod, while untreated wells were left as negative control. Fixation by 4% formaldehyde for 10 min, permeabilization by Triton x for 10 min and blocking by bovine serum albumin for 1 hour were done. Anti-TLR4 antibody (ThermoFisher; Cat. No. MA5-16216) and anti-TLR7 polyclonal antibody (Rabbit, anti-human, Abcam; Cat. No. ab45371) were added to the BCG and Imiquimod treated wells respectively on the coverslips. The plates were placed in the fridge overnight. Then cells were incubated with secondary antibodies Alexa fluor^®^555 goat anti-mouse (Life Technologies, Cat. No. A21422) and Alexa fluor^®^ 488 goat anti-rabbit (Thermofisher, Cat. No. A-11008) respectively for one hour followed by nuclear staining by Hoechst 33342: (Life Technologies, Cat. No. 62249) counter stain for one minute. Finally, coverslips were mounted on glass slides and examined by Confocal Microscope Leica TSC SPE II/ DMi 8 to visualize the cells and obtain images. Images were later analyzed using ImageJ 1.53e program to obtain the fluorescence intensity according to the following equation:

Corrected total cell fluorescence = Integrated density − (Area of selected cell × Mean fluorescence of background)

### Verification of TLR 2 Upon the Application of BCG by Flow Cytometry

Six-well plates of SCC-4 cells were prepared; 80,000 cells/well. When cells reached confluence, doses of BCG (0.5 and 5 mg/ml) were applied. Control wells were left untreated (Negative control). Trypsinization was then done followed by the transfer of the cell suspensions to falcon tubes. Falcon tubes were centrifuged at 1800 rpm for 5 min followed by fixation, permeabilization and staining with 1ry monoclonal anti-TLR2 antibody (ThermoFisher; Cat. No. MA5-16260) for 30 min at room temperature. This was followed by 2ry antibody in the dark for 1 hour. The cells were then transferred to flow cytometer tubes and analyzed by fluorescent-activated cell sorter FACScan flow cytometer (Becton Dickinson, equipped with Cell Quest Software, USA).

### Quantification of Autophagic Phenotype in SCC-4 cells

SSC-4 cells in T25 flasks were treated with BCG (0.05 mg/ml) and (0.1 mg/ml), Imiquimod (20 ug/ml) and (80 ug/ml), or Cisplatin (Unistin^®^) (Hikma, Egypt) (7 µM which equals 2µg/ml) for 30 min, 1 hour, 6 hours., 24 hrs., 48 hrs., 72 hrs., and 6 days. The indicated doses were selected to be sublethal (below IC50), based on the cell viability assays performed for BCG and imiquimod, or from previously published literature regarding cisplatin (an autophagy-inducing dose) [34,35]. At each definite time point, the cells were harvested by trypsinization then transferred to 15 ml falcon tubes and centrifugated at 1200 rpm for 5 min.

Then steps for staining with pre-conjugated LC3B antibody were carried out according to manufacturer’s guidelines. Cell suspensions were collected thoroughly in FACS tubes. Cell acquisition and data analysis of the stained cell suspensions was performed by FACS Calibur flow cytometer (Becton Dickinson, San Diego, Calif, USA) and Cell Quest software (Becton Dickinson) respectively. Samples were run in duplicates with about 10,000 events counted per tube. The negative control samples were run first to identify the position of the negative and the positive populations followed by the tubes containing the stained cell suspensions. During analysis of scatter plots, the percentages of cells undergoing autophagy were those in the upper right, and upper left quadrant collectively.

### Animal Experiments

A total of 60 male Syrian golden hamsters (*Mesocricetus auratus*), aged 5 weeks (80- 110 g), were obtained from Theodor Belhars Institute for Medical Research (Cairo, Egypt).

Animals were housed individually in showbox polypropylene cages (Technoplast, Italy) separately (one/cage) in the Medical Technology Center in the Medical Research Institute, Alexandria University. The hamsters underwent a two-week acclimation period prior to the start of the experiment. They were kept under controlled conditions with a room temperature of 23 ± 1 °C, relative humidity of 50 ± 5%, and a 12-hour light/dark cycle. Throughout the study, they had free access to drinking water and standard feed, provided in compliance with the guidelines of the Medical Research Institute. Animal house personnel cleaned the cages once per week.

### Ethics Statement for Animal Research

Experimental protocols were approved by the Alexandria University review committee (IRB#00010556-IORG0008839). All methods were carried out in accordance with the ARRIVE guidelines ([Home | ARRIVE Guidelines](https://arriveguidelines.org/)).

### Tumor Induction

We chemically induced OSCC by 7,12-dimethylbenz[a]anthracene (DMBA) (Sigma Aldrich, Cat No. D3254). We topically applied 0.5% DMBA dissolved in 100 ml liquid paraffin using number 4 paint brush to the left buccal pouch, with the right side left untreated to allow normal feeding. Carbamide peroxide was used as an accelerator, prepared by dissolving 64 gm of urea in 103 ml of hydrogen peroxide. Both DMBA and carbamide peroxide were administered alternately five days per week (three days DMBA, two days accelerator) to shorten tumor development period.[36,37] This protocol was followed until the development of intraoral or extraoral tissue changes indicative of SCC. Hamsters were examined and changes were recorded on a weekly basis.

### Grouping and Treatment Plan

Following tumor establishment, only 41 hamsters have survived. They were randomized into seven groups and were treated with the desired drug/drug combination according to the planned protocol. An insulin syringe was used for injections and a number 4 paint brush was used for applying Imiquimod cream.

BCG was injected intraperitoneally with a dose of 1 x 10^6^ colony forming units in 0.2 ml once weekly for six weeks [38]. Imiquimod was applied topically with a dose of 50 mg of the Aldara cream applied on the lesion with a safety margin of 0.5 cm, three times per week for 4 weeks [39]. It is important to note that although Imiquimod is applied topically, its systemic absorption is well documented in literature [40]. MPLA was injected intraperitoneally with a dose of 25 µg, once a week for 3 weeks [41]. Cisplatin was injected intraperitoneally with a dose of 7mg/kg body wt., once a week for 4 weeks [42]. The negative control group received no treatment. Group allocation, assessments, and analysis were blinded to investigators.

After treatment, the animals in each group were followed up for 2-4 weeks or until euthanization. Animals were euthanized post-treatment under anesthesia (ketamine 30 mg/kg, i.p.) by cervical decapitation. Carcasses were incinerated.

Lesion specimens were dissected and each one was divided into 3 pieces. The central piece for histological assessment. Another piece for autophagy detection by flow cytometry and the third for examination by transmission electron microscopy (TEM).

### Tumor volume reduction ratio and survival analysis

Tumor volume (TV) was measured before treatment by making two perpendicular measurements (D max: longer dimension; length) and (D min: shorter dimension; width) using a caliber **(Figure 3.6d)** and calculated as: (length*width^2^)/2

The same measurements were performed weekly during treatment and over a period of 4 weeks after treatment. The percentage of tumor volume change was calculated as:

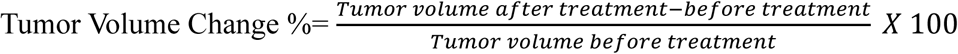

The animals’ survival rate was recorded and compared to the values of the control group. Increased life span (ILS %) was calculated as:

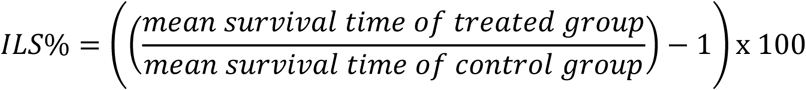

### Histopathological Assessment

After hamsters’ euthanization, the central part of the lesion specimen excised from each hamster was fixed in 10% neutral-buffered formalin solution for ∼24 hrs., to be processed for H&E staining and histopathological examination using light microscopy. Sections were blindly examined by 2 pathologists in randomly selected 5 microscopic fields at a magnification of ×400.

### Quantification of Autophagic Phenotype in Tumor cells

The second parts of the excised tumors were preserved in Roswell Park Memorial Institute (RPMI) 1640 media to be prepared for flow cytometry assessments for autophagy. The specimens were dissociated through mincing with sharp surgical blades in cold RPMI 1640 and trypsin/EDTA. The extracted cells were separated from the remaining tissue by filtration through 100 µm meshed cell strainer in a sterile 15 ml tube. Cells were centrifuged at 2000 rpm for 20 min., then incubated with trypsin for 20 min.

Cells were centrifuged again then washed by PBS and bovine serum albumin. Then, analysis of cell autophagy was done by staining with monodansylcadaverine (MDC) obtained from Autophagy Detection Kit (Abcam, ab139484) according to manufacturer’s guidelines. Finally, cells were analysed by FACScan.

### Assessment of Ultrastructural Morphology of Autophagic Tumoral Cells

We used TEM to identify autophagic cellular morphology. The third part of the fresh lesional specimens were trimmed into small pieces measuring 1 x 1 x 1 mm, fixed in 3% glutaraldehyde, washed in phosphate buffer, and then fixed in 1% osmium tetraoxide (OSO4) for 2 h at 4 °C. The specimens were dehydrated and finally embedded in araldite capsules. Sections were mounted on copper grids, double stained with freshly prepared uranyl acetate and lead citrate and finally examined using TEM (JOEL, JSM-6360LA, Japan).

### Statistical Analysis

In the *in vitro* study data are presented as mean ± SD. The significance of differences was evaluated through two-way ANOVA. The two-way ANOVA and Tukey’s multiple comparisons tests were used to analyze the differences between the test and control groups along the seven-time intervals. In addition, the differences between time intervals in each group was analyzed by the mentioned tests.

In the *in vivo* study, data are presented as mean ± SEM. One way ANOVA and Tukey’s multiple comparisons tests were used to analyze the differences in the tumor volume change and the increased life span percentages between groups. Kaplan-Meier method was used for survival analysis. Statistical significance was defined as a P-value less than 0.05 in all tests. The statistical analyses were performed with Graphpad Prism 9.0 (GraphPad Software, San Diego, CA, USA).

## Results

### OSCC Cells Respond to BCG and Imiquimod in a Dose-Dependent Manner but Are Insensitive to MPLA

Given that the goal of this study is to induce OSCC cells to commit to autophagy, dosages of TLR agonists needed to be determined. To ensure sublethal, autophagy-inducing conditions, we tested doses below the half -maximal inhibitory concentration (IC50). We calculated IC50 and IC80 for both BCG and Imiquimod (Fig. 1a,b).

**Figure 1.**
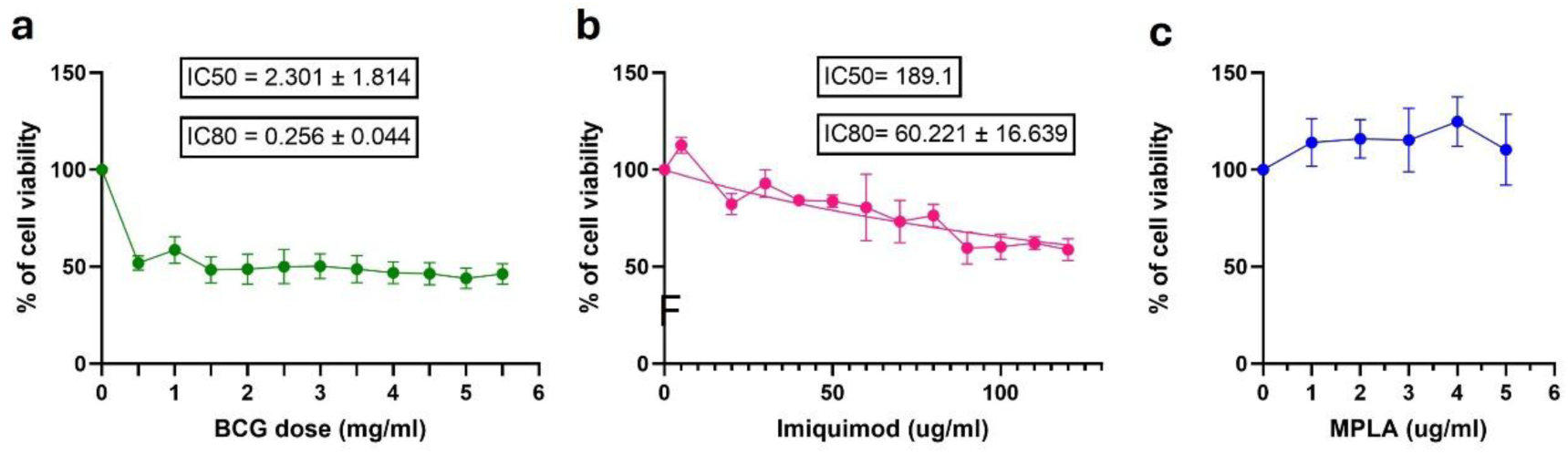
BCG and Imiquimod inhibited the growth of SCC4 cell line, while MPLA showed no cytotoxic effect. SCC4 cells were treated for 48 hr with increasing concentrations of (a) BCG, (b) imiquimod, or (c) MPLA. Cell viability was assessed by CCK-8 assay and normalized to untreated controls. Dose-response curves were plotted to calculate IC50 and IC80 values. Data are mean ± SD.

None of the MPLA doses revealed any cytotoxic effect, and the drug permitted SCC-4 cell proliferation (Fig. 1c). Based on the results of this test, MPLA was excluded from being further tested in the *in vitro* phase. Given its established role as a vaccine adjuvant that primarily acts through immune activation [43], MPLA was reserved for the in-vivo studies to test immune-mediated effects.

The next step in the current study was to validate TLRs activation in OSCC cell line in response to these agonists.

### BCG and imiquimod increase the expression of TLRs 2, 4, and 7 in OSCC cells

To determine whether BCG and imiquimod activate their target TLR pathways in OSCC, we examined TLR expression and localization in SCC-4 cells.

Confocal microscopy of TLR4-stained cells showed that BCG treatment shifted receptor distribution from predominantly membrane-associated to mainly nuclear, followed by cytoplasmic (Fig. 2a), suggesting altered receptor trafficking and non-canonical TLR4 signaling. Untreated cells exhibited aberrant localization to both nuclear and membrane-associated compartments. TLR7 staining was largely cytoplasmic, with stronger signal intensity in imiquimod-treated cells (Fig. 2b), consistent with ligand-induced receptor upregulation and activation within endosomal compartments. Quantitative analysis confirmed selective upregulation of TLR4 in response to BCG and TLR7 in response to imiquimod (Fig. 2c, d).

**Figure 2.**
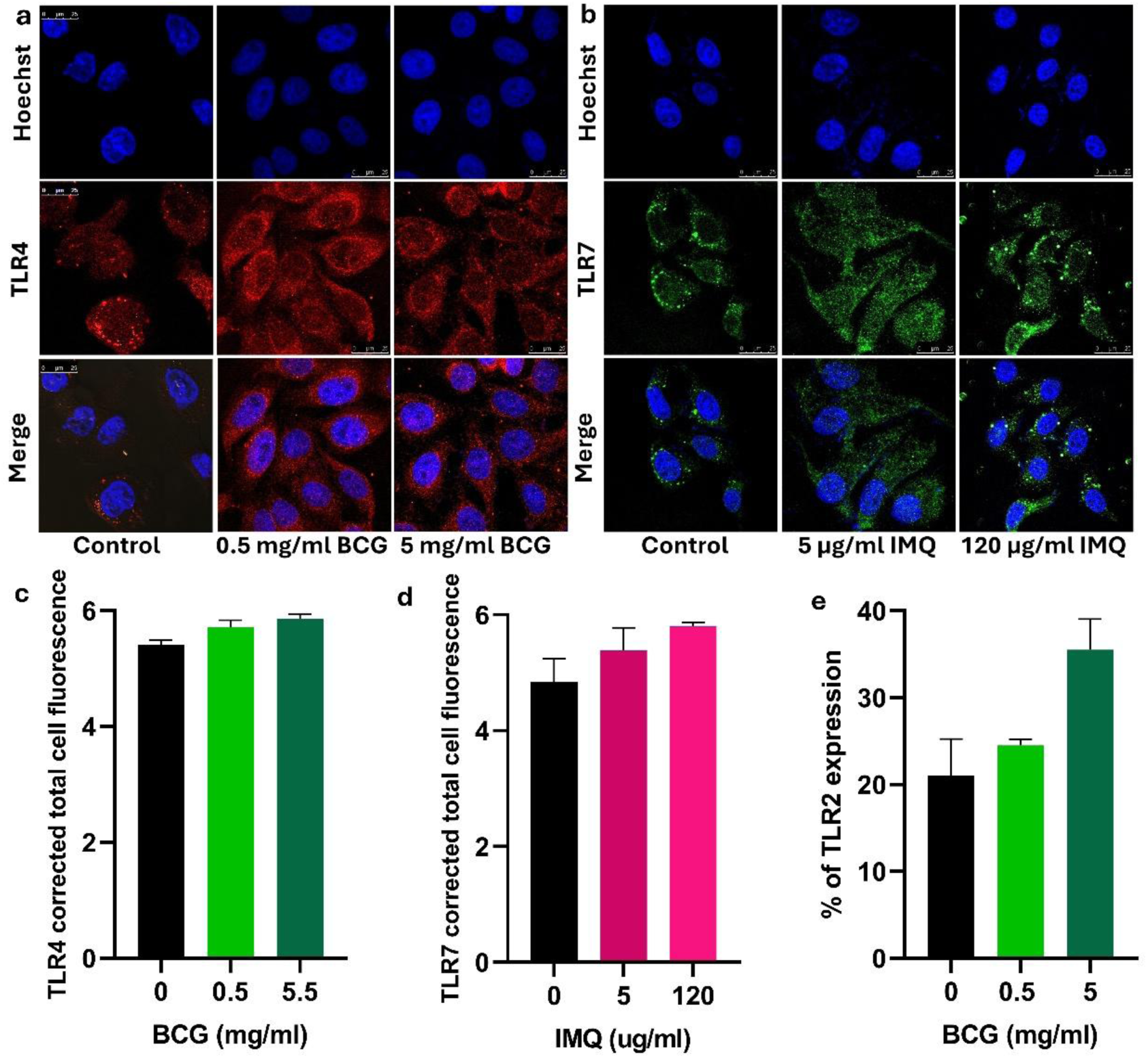
BCG and Imiquimod (IMQ) increase the expression of TLRs 2,4, and 7 in SCC-4 cells. SCC-4 cells were treated with BCG (0.5 and 5 mg/ml) or imiquimod (IMQ) (5 and 120 µg/ml) for 24 hr. (a-b) Confocal microscopy showing TLR4 and TLR7 expression. (c-d) Quantification of corrected total cell fluorescence. (e) Flow cytometry showing TLR2 upregulation after BCG treatment. Data are mean ± SD.

Flow cytometry further revealed increased TLR2 expression following BCG treatment (Fig. 2e), indicating broader TLR pathway engagement, likely via cross-talk with TLR4. These findings confirm target engagement for both TLR agonists and raise the question of whether such activation leads to downstream functional effects, such as autophagy induction.

### BCG (TLR2/4 agonist) induces autophagy in OSCC cells

To determine whether TLR2/4 activation triggers autophagy in OSCC, SCC-4 cells were treated with sublethal doses of BCG (0.05 mg/ml and 0.1 mg/ml) and monitored for LC3B expression at multiple time points. Scatter plots are found in supplementary fig. 1.

Both BCG doses produced a biphasic autophagy response (Fig. 3a). The percentage of autophagic cells increased up to 6 hr, dropped at 24 hr, and peaked again at day 6.

**Figure 3.**
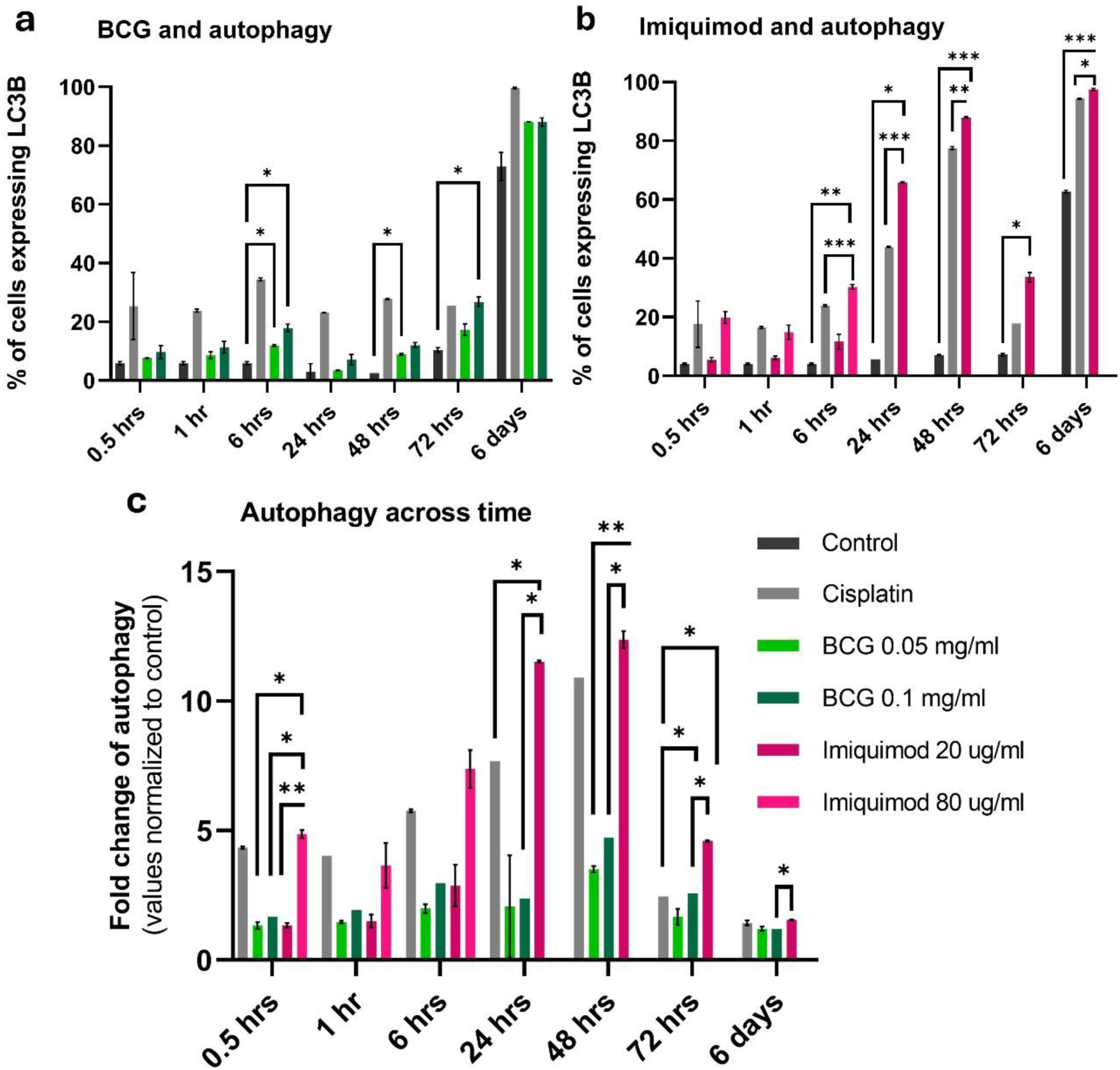
Effects of TLR agonists BCG and Imiquimod on autophagy compared to Cisplatin. (a-b) Time course of autophagy induction measured by LC3B staining after treatment with BCG (0.05 and 0.1 mg/ml), imiquimod (20 and 80 µg/ml), or cisplatin (2 µg/ml). Autophagy was quantified by flow cytometry across seven time points (0.5-144 hr). (c) Fold-change in autophagy relative to untreated controls, showing imiquimod as the strongest inducer compared with BCG or cisplatin. Data are mean ± SD. Two-way ANOVA with Tukey’s test; *P < 0.05, **P < 0.01, ***P < 0.001, ****P < 0.0001.

Cisplatin and untreated controls also showed elevated autophagy at day 6, suggesting that late-stage increases may reflect general stress rather than a drug-specific effect. At earlier intervals, BCG induced significantly greater autophagy than controls (0.05 mg/ml at 6 hr and 48 hr; 0.1 mg/ml at 6 hr and 72 hr). The overall similarity between the 2 doses suggests autophagy induction was dose-independent within this range.

These results show that BCG rapidly triggers autophagy in OSCC cells, prompting the question of whether similar effects occur following TLR7 activation by imiquimod.

### Imiquimod (TLR7 agonist) induces autophagy in OSCC cells

To determine whether TLR7 activation triggers autophagy in OSCC, SCC-4 cells were treated with sublethal doses of imiquimod (20 µg/ml and 80 µg/ml) and assessed for LC3B expression over time. At 80 µg/ml, imiquimod formed an insoluble precipitate beyond 6 hr, interfering with LC3B staining; therefore, only early time points (30 min, 1 hr, and 6 hr) were evaluated for this dose. Scatter plots are found in supplementary fig. 1.

Autophagic cell percentages were higher in all imiquimod-treated groups compared to the negative control at every tested interval. At the three early time points tested for both doses, 80 µg/ml yielded higher autophagy levels than 20 µg/ml, but differences were not statistically significant (Fig. 3b), mirroring the pattern observed with BCG. This indicates that imiquimod-induced autophagy in OSCC cells is largely independent of the tested concentration range.

Given that both TLR agonists induce autophagy, we next compared their effects directly with cisplatin to determine relative potency.

### BCG and imiquimod are comparable to cisplatin in inducing autophagy in OSCC cells

To compare the level of autophagy induced by TLR agonists with that induced by Cisplatin (2 µg/ml) (a gold standard chemotherapeutic for solid tumors and a known autophagy inducer in OSCC [34,35]), we analyzed data from parallel experiments (Fig. 3a,b). Cisplatin consistently induced autophagy at all tested time points, with relatively stable levels across the time course, a slight increase at 6 hr, and a peak at day 6. Scatter plots are found in supplementary fig. 1.

LC3B expression in the cisplatin group was significantly higher than in BCG-treated cells at all time points except 72 hr. In contrast, imiquimod induced significantly greater autophagy than cisplatin at most later intervals (24 hr to 6 days), indicating a sustained and potent autophagic inducer, surpassing even a classical chemotherapeutic agent.

Overall, imiquimod emerged as the most potent and sustained inducer of autophagy, exceeding cisplatin at later time points, whereas BCG produced a steady but comparatively moderate effect. Strikingly, despite cisplatin’s established potency as a cytotoxic and autophagy-inducing drug, both BCG and especially imiquimod matched or surpassed its autophagic effect. This suggests that TLR activation can provoke robust tumor-intrinsic stress responses through mechanisms distinct from DNA damage. These observations raised the question of whether the high autophagy levels observed reflect drug-specific activity or are partly due to general increases in autophagy over time.

### Fold-change analysis reveals a strong association between the TLR7 agonist imiquimod and autophagy induction in OSCC

To distinguish drug-specific effects from time-dependent autophagy increases in untreated cells, as well as negate basal autophagy levels, we normalized autophagy values to the corresponding negative controls (Fig. 3c). For comparison, cisplatin values from two independent experiments—BCG and imiquimod—were averaged after normalization.

After normalization, BCG induced lower autophagy than cisplatin at all time points except 72 hr, with no differences between doses. The pronounced day-6 spike seen in the raw data disappeared, indicating it reflected a general, time and stress-dependent increase rather than a BCG-specific effect.

In contrast, imiquimod remained the most prominent inducer after normalization. The 80 µg/ml dose produced the highest induction at 0.5 and 6 hr, whereas the 20 µg/ml dose rose sharply at 24 hr and peaked at 48 hr before declining by day 6.

Despite differences in magnitude, all agents showed a similar temporal pattern, early induction, mid-phase fluctuation, and late decline, suggesting a shared response dynamic. Among them, imiquimod generated the strongest drug-specific autophagic effect. This finding led us to investigate whether these *in vitro* autophagy patterns translate into functional anti-tumor effects *in vivo*.

### BCG, imiquimod, and their combinations with MPLA exert potent *in vivo* antitumor activity

We compared cisplatin, the standard OSCC chemotherapeutic, with TLR agonists BCG (TLR2/4) and imiquimod (TLR7), alone or in combination, as well as MPLA as an adjuvant, following the treatment schema in Fig. 4. To our knowledge, this is the first in vivo study to directly evaluate these agents and combinations within a single OSCC model. The experiment included 41 Syrian golden male hamsters with DMBA-induced OSCC in the left buccal pouch. Representative pre- and post-treatment images are shown in Fig. 5a. Tumor volume (Fig. 5b) and survival (Fig. 5c,d) were the primary clinical endpoints.

**Figure 4.**
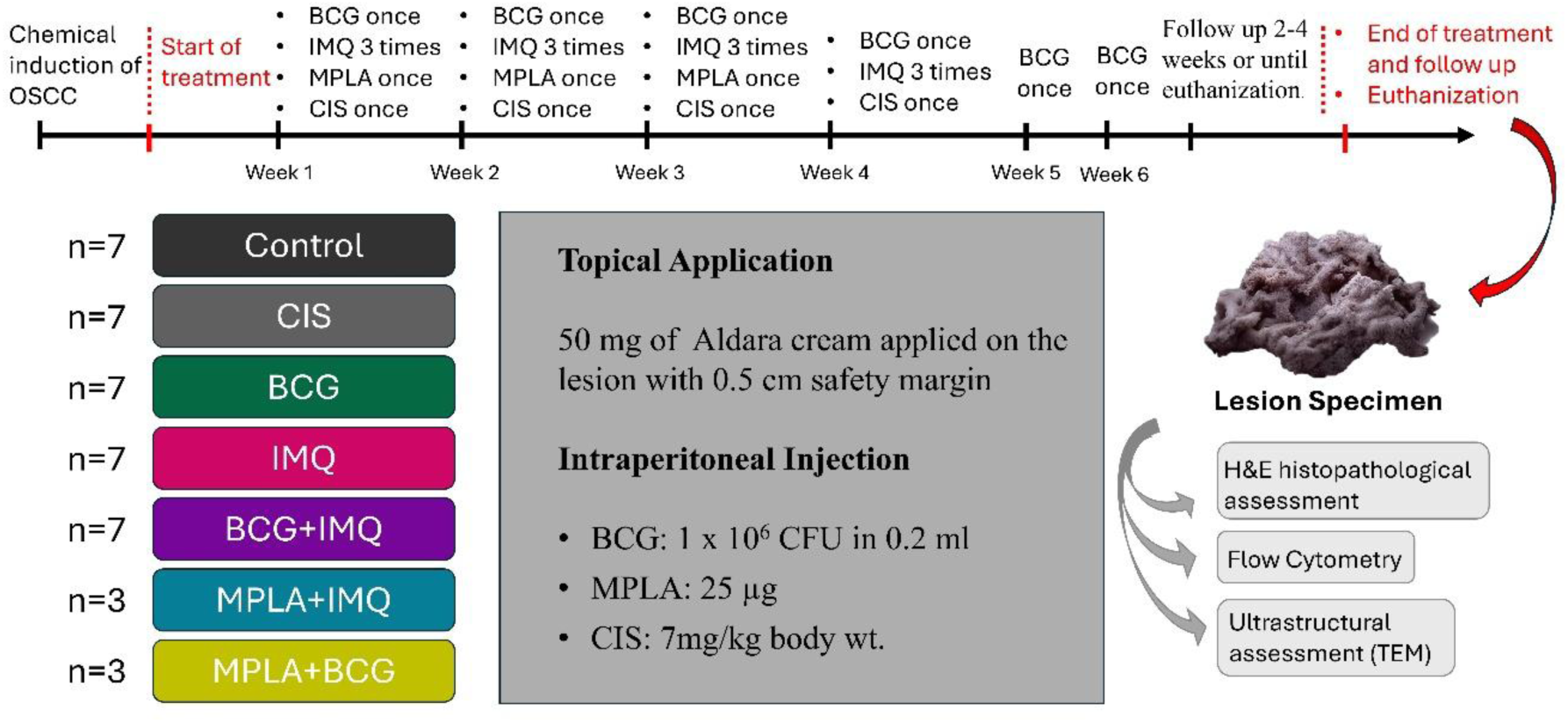
*In vivo* treatment schema. Treatment groups included: Control, Cisplatin (7 mg/kg), BCG (1 × 10^6 CFU), Imiquimod (IMQ) (50 mg topical), BCG+IMQ, MPLA+IMQ, and MPLA+BCG. Regimens are illustrated in the schema. Downstream analyses included histopathology, flow cytometry, and TEM.

**Figure 5.**
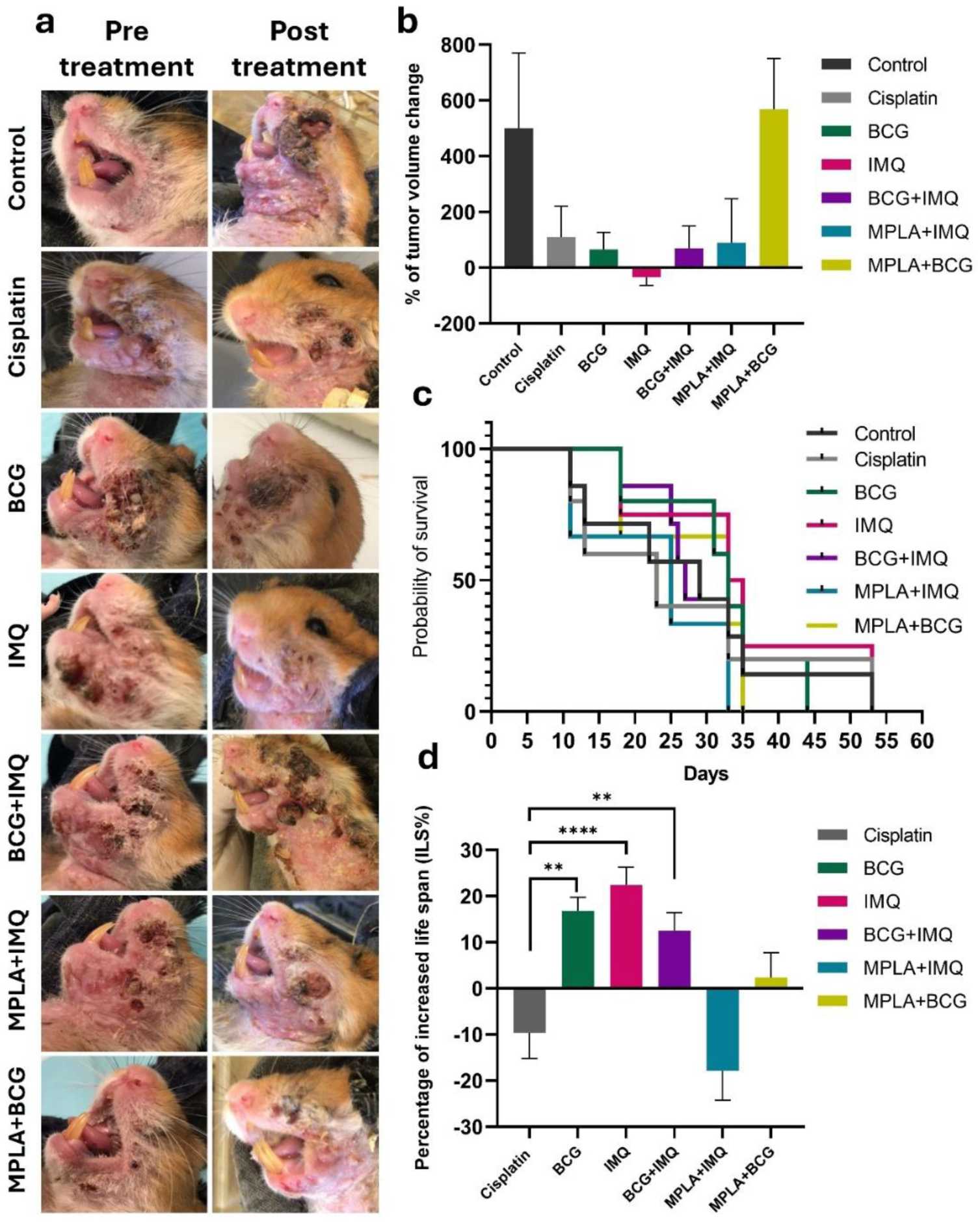
BCG, Imiquimod (IMQ), and their combination with each other as well as with MPLA exert a remarkable *in-vivo* antitumor activity. **(a)** Oral squamous cell carcinoma lesions pre and post treatment. Tumors presented as exophytic masses, with evident areas of ulceration and necrosis. The post-treatment images were taken after completion of each treatment (either after 4 weeks or 6 weeks). (b) Percentage of tumor volume change. A decrease in the tumor volume occurred only after treating the hamsters with Imiquimod. In BCG, BCG+IMQ combination, and MPLA- IMQ combination treated groups, the % of increase in tumor volume was less than Cisplatin. (c) Kaplan Meier Survival Curve for Treatment Groups. The survival analysis was plotted after recording the death dates until the end of the experiment on day 53. (d) Percentage of Increased Life Span (ILS%). BCG, Imiquimod, and their combination showed a significant increase in ILS% of hamsters in comparison to Cisplatin which revealed a significant decrease in ILS%. *P < 0.05, **P < 0.01, ***P < 0.001, ****P < 0.0001. Data are expressed as mean ± SEM.

In untreated controls, tumor volume increased by ∼500%, reflecting rapid disease progression. Imiquimod was the only treatment to achieve an actual tumor volume reduction (>30%), demonstrating strong antitumor activity. BCG, BCG-imiquimod, and MPLA-imiquimod combinations also achieved better tumor control than cisplatin, whereas the MPLA-BCG combination performed worse than the negative control.

Survival analysis revealed that BCG, imiquimod, and their combination significantly prolonged lifespan compared with cisplatin, which paradoxically reduced survival by ∼10%. These findings highlight the advantage of TLR agonists over standard chemotherapy: effective tumor control with lower morbidity and mortality. Based on the strong in vitro evidence linking these agents to autophagy induction, we next assessed whether autophagy contributed to their antitumor effects in vivo.

### BCG, imiquimod, and their combinations induce autophagy *in vivo*

Given our in vitro findings that BCG and imiquimod strongly induce autophagy in OSCC cells, we next investigated whether this mechanism also occurs in vivo. Demonstrating autophagy induction in tumor tissue following TLR agonist treatment would strengthen the mechanistic link between TLR activation and tumor control in our animal model.

Histopathological analysis of tumor sections revealed regions with extensive cytoplasmic vacuolization, particularly in highly cellular tumor areas near lymphovascular invasion (Fig. 6a,b). Although autophagy cannot be directly visualized by light microscopy, these features are compatible with autophagic processes and may represent structural changes associated with cell detachment.

**Figure 6.**
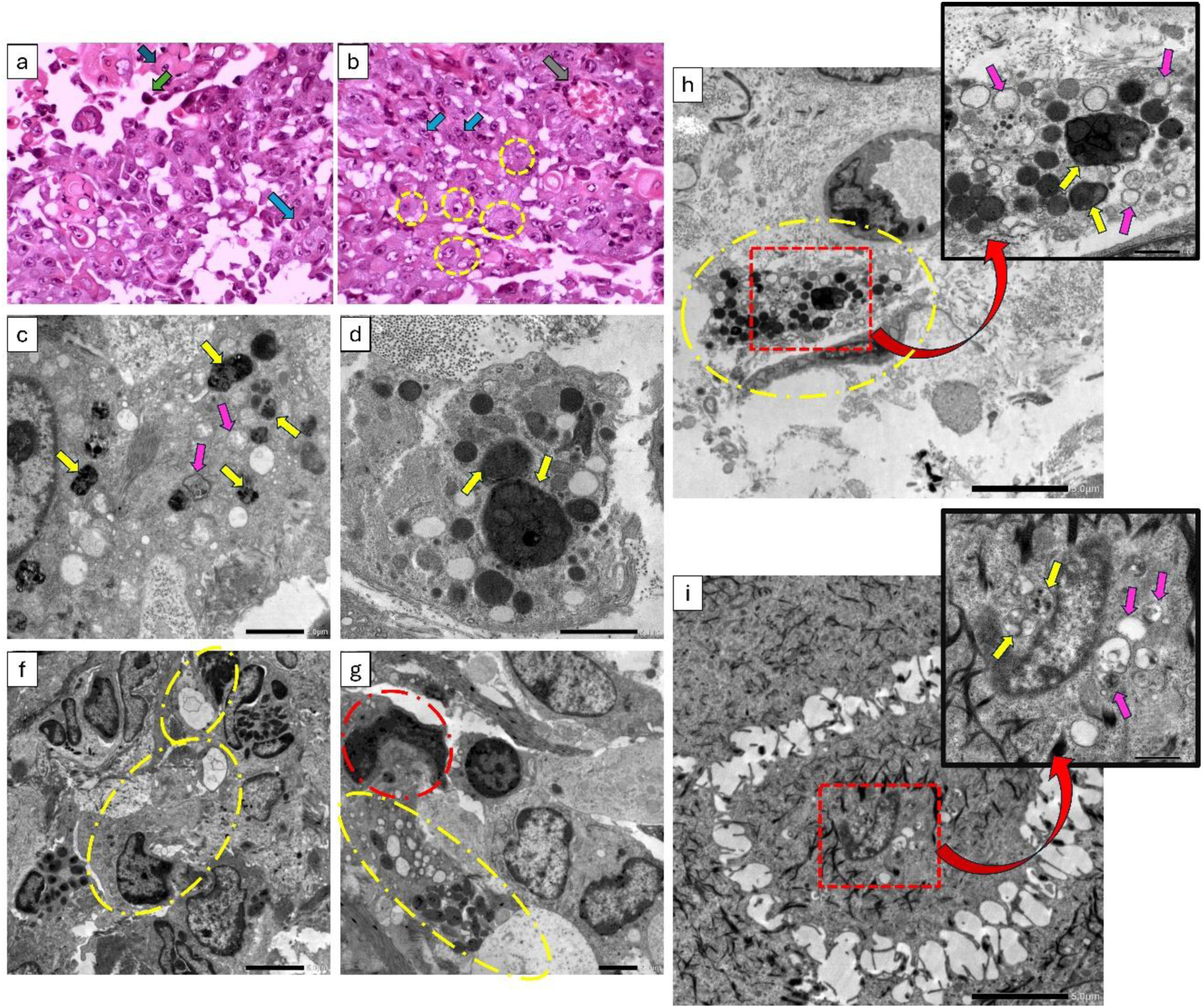
Histopathological and ultrastructural evidence of autophagy. (a-b) H&E-stained sections (X400) from the BCG-treated group show anaplastic epithelial cells with loss of desmosomal attachments, intra-epithelial cleft formation, mitotic figures (yellow arrows), individual cell keratinization (blue arrows), apoptotic cells (green arrows), and extensive vacuolization suggestive of autophagy (yellow circles). Grey arrow indicates lymphovascular invasion. (c-g) TEM images from the imiquimod-treated group illustrate autophagosomes (pink arrows) and autolysosomes (yellow arrows, irregular single-membrane structures). Cells with extensive vacuolization but no chromatin condensation consistent with ADCD are shown (f, yellow circles). Panel (g) highlights the contrast between ADCD (lower cell) and apoptosis (upper cell, red circle). (h-i) TEM images from the IMQ+BCG combination group. (h) shows a cell undergoing ADCD with extensive vacuolization, while (i) depicts an apoptotic cell with nuclear shrinkage and chromatin disintegration occurring alongside autophagy. Insets show higher magnification of autophagic vacuoles.

Ultrastructural examination by TEM confirmed hallmark features of autophagy. Autophagosomes with double membranes and autolysosomes with single irregular membranes containing degraded material were observed [44] (Fig. 6c,d). Cells with extreme vacuolization but without chromatin condensation, consistent with ADCD, were also noted (Fig. 6f-h) [18,45] In contrast, apoptotic cells exhibited nuclear shrinkage, chromatin condensation, and membrane blebbing, sometimes occurring alongside autophagic vacuoles (Fig. 6g,i) [18,46]

Quantitative analysis by flow cytometry further confirmed that all treatment groups induced significantly higher autophagy levels than control (Fig. 7). Tissue yield was insufficient for the cisplatin and MPLA-imiquimod groups due to poor survival in these animals, precluding analysis in those arms. The BCG-imiquimod combination induced the strongest response, suggesting synergy between TLR2/4 and TLR7 activation.

**Figure 7.**
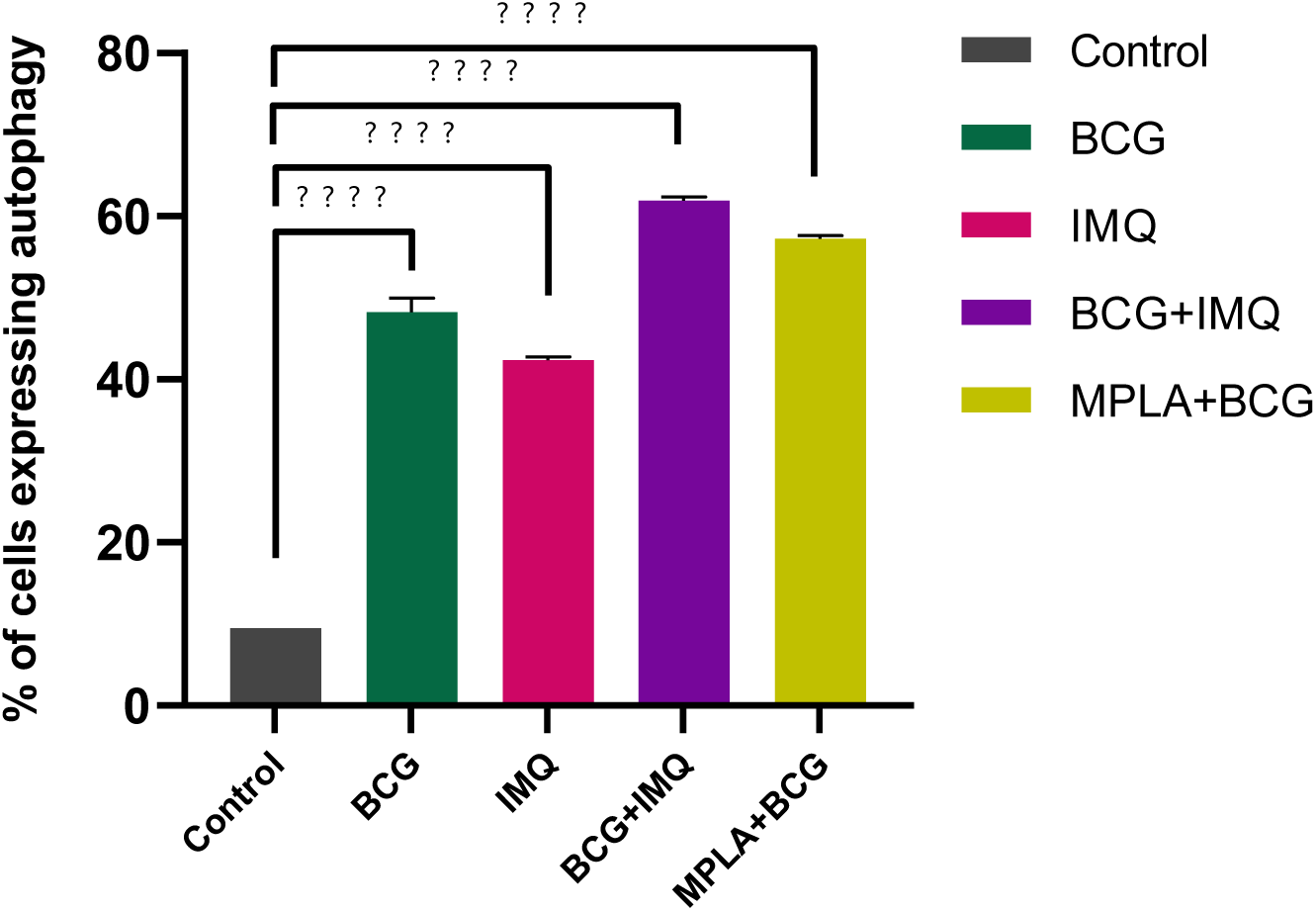
Expression of Autophagy in Treatment Groups *In-vivo*. Flow cytometry analysis of tumor specimens revealed significantly higher percentages of autophagic cells in BCG, imiquimod, BCG-imiquimod, and MPLA-BCG groups compared to untreated controls. *P < 0.05, **P < 0.01, ***P < 0.001, \*\*\**P < 0.0001. Data represent mean ± SEM*.

MPLA-BCG also exceeded BCG alone, reinforcing the association between TLR signaling and autophagy in vivo. Interestingly, imiquimod alone induced lower autophagy than BCG, in contrast to the in vitro results, suggesting that additional *in vivo* factors modulate the magnitude of autophagy.

Together, these findings indicate that the most effective regimens for tumor control, particularly BCG and imiquimod, are also those that trigger robust autophagy in vivo, supporting a potential mechanistic role for autophagy in their therapeutic efficacy.

## Discussion

To our knowledge, this is the first study to directly link the antitumor effects of the FDA-approved [47] TLR agonists BCG and imiquimod in OSCC with autophagy modulation, integrating both mechanistic and therapeutic endpoints.

BCG has been investigated in head and neck cancers since the 1970s, but earlier trials were limited by heterogeneous design and inconsistent outcomes [21–26]. In our model, BCG reduced tumor volume more effectively than cisplatin and significantly prolonged survival, highlighting its favorable therapeutic profile with potentially fewer side effects than conventional chemotherapy. Imiquimod produced the most pronounced tumor regression, with more than a 30% volume reduction and survival benefit, consistent with clinical reports in cutaneous SCC, case reports of oral lesions and a recent single-arm pilot clinical trial [27–32]. Our findings suggest that the oral mucosa’s architecture may enhance drug penetration, supporting its activity even at lower doses.

Combination strategies yielded mixed results. While BCG-imiquimod improved outcomes relative to controls and cisplatin, it was less effective than either drug alone, likely reflecting the complexity of TLR crosstalk. Excessive or dysregulated signaling may shift from tumor suppression to immune evasion or stress adaptation. Similarly, MPLA-imiquimod improved tumor control but shortened survival, and MPLA-BCG unexpectedly increased tumor volume yet still extended survival, underscoring the context-dependent nature of TLR biology.

Mechanistically, both BCG and imiquimod induced autophagy *in vitro* and *in vivo*. In untreated controls, low baseline autophagy likely reflects early suppression, a known tumor-initiating mechanism [48–50]. In treated groups, autophagy appeared to act as a therapeutic effector, correlating with tumor regression and survival benefit, especially in single-agent regimens. Notably, very high autophagy levels in some combinations coincided with poorer outcomes, reinforcing that while autophagy can promote tumor cell death, excessive or prolonged activation may instead support survival under stress. The concept of autophagy-dependent cell death (ADCD) remains debated [18,51]; our data suggest its contribution but cannot prove it without genetic or pharmacological validation, which should be addressed in future studies.

On a side note, we uncovered an altered pattern of TLR4 expression in both BCG and control groups that suggests a deviation in receptor trafficking in OSCC [52] and non-canonical TLR4 signaling. This provides a plausible factor regulating response to TLR therapy and encourages further research.

An additional finding was that cisplatin induced autophagy in OSCC cells, challenging the traditional view that autophagy uniformly drives cisplatin resistance. This aligns with emerging evidence that cisplatin’s efficacy may partly rely on autophagic mechanisms [53,54], warranting further exploration.

Together, these findings position BCG and imiquimod as promising immunotherapeutic candidates for OSCC. Their efficacy appears partly mediated through TLR-driven autophagy, though outcomes depend strongly on context and dosing. Careful optimization of single-agent versus combination strategies, along with deeper mechanistic studies, will be essential to translate these insights into effective clinical interventions.

## Conclusions

Our work emphasizes the promise of imiquimod and BCG as immunotherapeutic agents in the context of OSCC. Our framework integrating *in vivo* and mechanistic studies offers a novel perspective on the role of autophagy-mediated cancer cell death in cancer treatment, that also challenges the prevailing notion that autophagy drives cisplatin resistance. These findings warrant further investigation into these immunotherapeutics and the complex role of autophagy in cancer progression. Importantly, this research provides a strong translational basis for developing more effective, autophagy-targeted immunotherapeutic strategies for OSCC patients.

## Supporting information

This document contains additional information regarding the materials and methods section, in addition to an additional figure.

## Abbreviations

ADCD: Autophagy-dependent cell death
ATCC: American Type Culture Collection
BCG: Bacillus Calmette-Guérin
CCK-8: Cell counting kit-8
CERRMA: Center of Excellence for Research in Regenerative Medicine and its Applications
DMBA: 7,12-dimethylbenz[a]anthracene
IC50: Half -maximal inhibitory concentration
ILS: Increased life span
MPLA: Monophosphoryl-lipid A
OSCC: Oral squamous cell carcinoma
RPMI: Roswell Park Memorial Institute
TEM: Transmission electron microscopy
TLRs: Toll-like receptors

## Declarations Ethics Approval

Experimental protocols were approved by the Alexandria University review committee (IRB#00010556-IORG0008839). All methods were carried out in accordance with the ARRIVE guidelines (<u>Home | ARRIVE Guidelines</u>).

**Clinical Trial Number** Not applicable

**Consent to Participate** Not applicable

## Consent for Publication

Not applicable.

## Data Availability

The data generated and used during the current study are available from the corresponding author on reasonable request.

## Authors’ Contributions

M.M.A. and E.M.O. conceived the project, designed the study and obtained funding.

M.M.A supervised the research, validated the findings, and contributed to reviewing and editing the manuscript. R.A.M and M.I.E supervised the research and validated the findings. N.A.S. and M.M.A wrote the manuscript. N.A.S., M.N.M., A.A.E., conducted experiments and analyzed the data. H.M.A., O.R.M. and E.M.O. conducted experiments and assisted with study design. R.A.M and A.K.A. performed flow cytometric analysis and assisted with experiment design.

## Acknowledgements

The cell line (SCC-4) used in this research was a generous gift from Dr. Rania Younis, University of Maryland, USA. We thank Dr. Marwa M. Essawy for her precious insights and generous advice during data analysis.

