## Supplementary material for "Autophagy-Mediated Antitumor Effects of BCG and Imiquimod in Oral Squamous Cell Carcinoma": This document contains additional information regarding the materials and methods section, in addition to an additional figure.

#### Material and Methods

##### Drugs Preparation

BCG was provided as 30mg/ml vial in a liquid form that did not need any special preparation. Imiquimod was provided as 200mg powder vial, soluble in Dimethyl sulfoxide (DMSO) and poorly soluble in water. The powder was dissolved in a mixture of 10 µl of DMSO and 990 µl of CCM, then a water bath sonicator (high frequency vibrator $20x10^6$ / sec) was used to help in the solubilization of the drug. The MPLA vial -100 µg ready-made solution with 0.5 mg/ ml concentration (500 ug /ml)- did not need any special preparation.

### Figure and figure legend

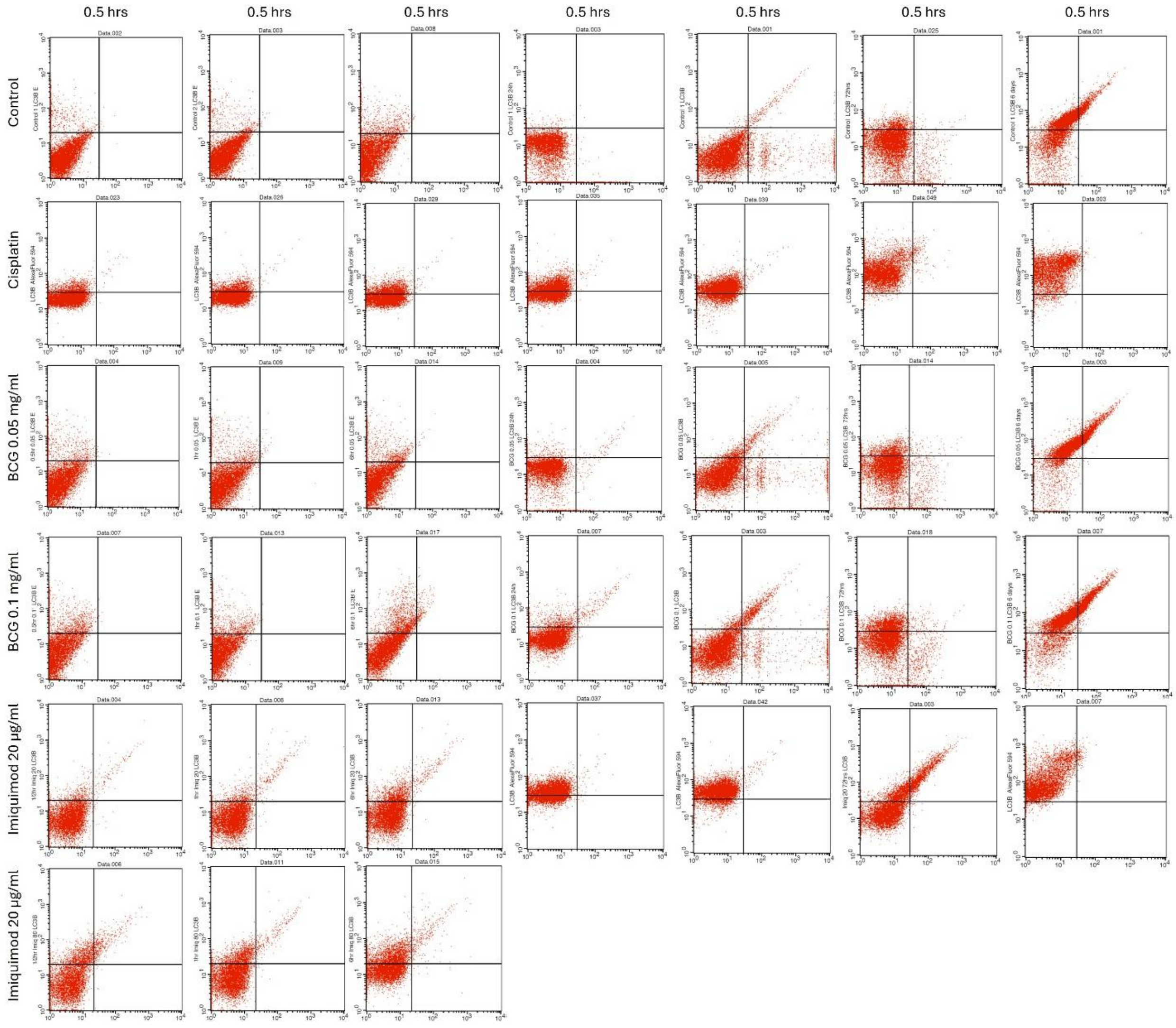


**Supplementary Figure 1. Scatter Plots for Human LC3B Alexa Fluor® 594-conjugated Antibody using Flow Cytometry at 7 Different Time Points.** 30 min., 1 hr., 6 hrs., 24 hrs., 48 hrs., 72 hrs., and 6 days in untreated SCC-4 cells and cells treated with cisplatin, BCG 0.05 mg/ml, BCG 0.1 mg/ml, Imiquimod 20 µg/ml and Imiquimod 80 µg/ml (only the first 3 time points are available for Imiquimod 80 µg/ml). The percentages of cells undergoing autophagy are presented in the upper right, and upper left quadrant collectively.
